# Path Integration Promotes Flexible Navigational Decision Making

**DOI:** 10.64898/2026.10.01.756060

**Authors:** Ryan Grgurich, Shiyun Wang, Yiting Wang, Jenna Pimenta, Kimia Delafraz, Hugh T. Blair

## Abstract

Flexible decision making depends on internal representations that capture the latent structure of a task, allowing prior knowledge to be applied in novel circumstances. Spatial navigation provides a useful model for studying flexible choice behavior, because spatial location is itself a latent variable that must be inferred rather than directly perceived. Here, we trained rats on a navigation task that required them to reach the same goal via two distinct paths from different starting locations, and then return to one of the two starting locations before beginning the next trial. Some rats were trained to rely on distal visual landmarks, allowing task states to be identified from current sensory information, whereas others were trained to rely on path integration, requiring task states to be inferred from self-motion information accumulated over the preceding trajectory. Rats trained to rely on path integration acquired the task similarly to rats trained to rely on distal visual landmarks. However, reliance on path integration enhanced rats’ ability to generalize their learning to novel situations, including finding the goal from an untrained starting location and adapting to a relocated goal that reversed learned choice sequences. These findings suggest that reliance on path integration promotes the acquisition and use of mental representations that capture the latent spatial structure of navigational tasks, thereby favoring flexible navigational decisions over rigid retrieval of reinforced responses. More broadly, our findings demonstrate how information available during training can shape representations learned from experience and thereby constrain the flexibility of decision making during inference.

## INTRODUCTION

Flexible problem solving depends on internal representations that capture the latent structure of a problem domain to support prediction, inference, and generalization (Gershman & Niv, 2010; Bengio et al., 2013). Artificial and biological neural networks can construct such representations by discovering structure in data, allowing past experience to guide decisions in novel situations (Radulescu et al., 2021; Assran et al., 2023; Li et al., 2024; Hafner et al., 2025). However, inferring latent structure from prior experience is constrained by the information available in the training data: different observations of the same underlying problem domain may produce representations with different capacities for inference.

Spatial navigation provides a useful model for investigating this principle because variables such as allocentric position and direction are themselves latent task dimensions that must be inferred rather than directly perceived. A rich literature of animal navigation studies has explored how wayfinding problems can be solved using different representations and strategies. “Route-based” representations can associate environmental cues with reinforced actions, supporting efficient travel along familiar routes but limiting flexibility when starting locations, routes, or goals change (Hull, 1943; Spence & Lippitt, 1946). In contrast, “map-based” representations capture latent spatial relationships among locations, allowing acquired knowledge to support inference when novel wayfinding problems arise (Tolman et al., 1946; Tolman, 1948; O’Keefe & Nadel, 1978). Even simple navigational decisions, such as choosing to turn left or right at a T-intersection, can be solved using different strategies (Ritchie et al., 1950; Hicks, 1964) that depend upon different brain regions (Packard & McGaugh, 1996; Packard & Goodman, 2013).

If the same navigational problem can be solved in different ways, then what determines which strategy an animal uses to find its way to a goal? Navigation strategies vary systematically with learning history, environmental cues, and cognitive demands (Blodgett & McCutchan, 1948; Restle, 1957; Gardner et al., 2013; Goodman, 2021), suggesting that an important determinant of navigational decisions may be the information represented in an animal’s current “task state,” the internal representation of its current situation used to select an appropriate action. Task states can differ in their dependence on temporal context: they may be identified from current sensory observations alone, such as familiar configurations of spatial landmarks (Knierim & Hamilton, 2011), or depend upon the sequence of observations and actions through which they were reached. They can also differ in whether they incorporate latent spatial variables such as allocentric position and direction. Path integration provides one means of inferring such variables by using self-motion signals accumulated along a trajectory to estimate displacement from a previously known state (Etienne & Jeffery, 2004).

It has been proposed that reliance on path integration to infer task states promotes cognitive-map formation and flexible navigation (Gallistel, 1990; McNaughton et al., 1996, 2006; Etienne et al., 1998). Computational models suggest possible mechanisms for this process by demonstrating how structured spatial representations can emerge through predictive and relational learning across sequential experience (Whittington et al., 2020; Stöckl et al., 2024; Gornet & Thomson, 2024). However, testing this hypothesis empirically is challenging, as it requires experimentally decoupling path integration from other sources of task state information, such as external spatial cues (Etienne et al., 1988). While such methods have successfully illuminated path integration’s contributions to self-localization, homing, and neural representations of space (Mittelstaedt & Mittelstaedt, 1982; Blair & Sharp, 1996; Bures et al., 1997, 1998; McNaughton et al., 2006; Jayakumar et al., 2019), less work has tested the prediction that reliance on path integration during navigational learning promotes flexible decision making during subsequent inference under changed routes or goal contingencies.

Here we addressed this question using the two-start two-choice (2S2C) task, which independently manipulated the informativeness of path integration and distal visual landmarks while rats learned the same navigational problem. We then tested their ability to navigate from an untrained starting location and adapt to a relocated goal. Reliance on path integration promoted flexible navigational inference consistent with map-based representations, whereas forced reliance on distal landmarks favored more rigid, route-based strategies. Rats trained with both information sources exhibited distinct navigational profiles, indicating that different strategies can emerge under identical conditions. These findings address how informational demands during learning shape acquired representations and the behavioral strategies they support, thereby influencing flexible decision making.

## RESULTS

The 2S2C task was performed on a plus maze embedded within a 2 × 2 m square track (Fig. 1A, B, C). Automatic doors controlled access to different maze segments. Four identical video monitors were positioned 1 m above the sides of the maze and were dimly backlit against surrounding black curtains (Fig. 1B), making them visible from most locations on the maze and forming a symmetric configuration of distal visual cues. In visually cued versions of the task, a distinctive visual landmark (a white triangle) was displayed on one of the four monitors, breaking the rotational symmetry of the array and providing directional landmark information. Fluid dispensers for delivering reward (100 μl sweetened condensed milk) were located at each of the four maze corners (Fig. 1B). Reward was delivered when rats entered the rewarded corner. Decoy vials of sweetened condensed milk were placed beneath all four corners to prevent olfactory guidance to the rewarded location. Track segments were rearranged into a new random spatial configuration each day to prevent rats from using uncontrolled local maze features as navigational cues.

**Figure 1.**
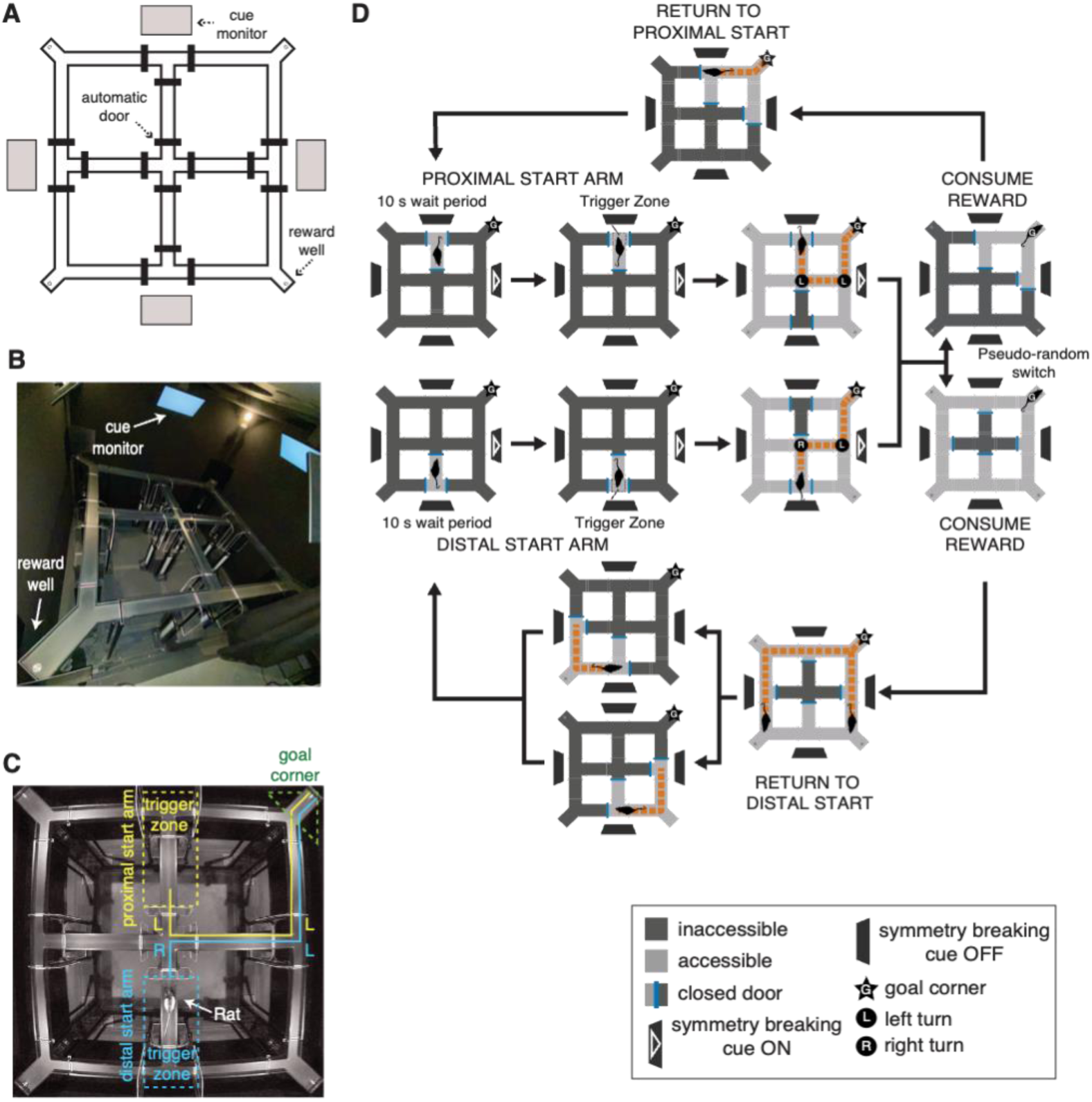
Maze apparatus and 2S2C task structure. A) Schematic of maze layout: small circles mark reward wells in the corners, heavy black lines mark automatic doors, and gray rectangles mark overhead monitors. B) Side view of the maze environment. C) Overhead view of a rat confined to the distal start arm at the beginning of a trial; yellow and blue lines show paths to the goal corner from the proximal and distal start arms, respectively. D) Flow chart of the 2S2C task structure: each trial starts from the proximal (top) or distal (bottom) start arm and ends when the rat reaches the goal corner; the automatic doors are then opened to direct the rat back to one of the two start arms to begin the next trial.

### Acquisition Training

Male and female Long-Evans rats were trained in daily sessions to obtain reward by navigating to a fixed goal corner of the square track (Fig. 1C). Each rat was assigned a goal corner (NW, NE, SE, or SW) within the task reference frame, counterbalanced across rats, and trained to reach that corner from two interior start arms within the plus maze: one proximal and the other distal to the goal. Each trial began from one of these two start arms. The task reference frame (including the start arms, goal corner, and visual landmark when present) was rotated each day with respect to the room reference frame to prevent rats from using uncontrolled distal room cues for navigation.

Maze doors were configured so that, from either starting location, the optimal path to the goal required passing through two sequential choice points: the 1st at the center of the maze and the 2nd at its perimeter (Fig. 1D). At each choice point, rats could turn either left (L) or right (R). From the proximal start arm, the optimal route required repeating the same turning response at both choice points (either LL or RR, counterbalanced across rats), whereas from the distal start arm, the optimal route required alternating turns (either LR or RL). Thus, the correct response at the 1st choice point depended on the starting location, whereas the correct response at the 2nd choice point was always the same.

Rats received reward upon reaching the goal corner at the end of each trial, regardless of whether they arrived via the optimal two-choice route or a longer, non-optimal route. While reward was consumed, maze doors were reconfigured to guide the rat back to a pseudorandomly selected proximal or distal start arm for the next trial. In visually cued variants of the task, the distal landmark cue was turned off as the rat traveled from the goal corner to the next start arm, leaving only the rotationally symmetric configuration of four backlit monitors visible during the return trajectory (Fig. 1D; Video S1).

#### Experimental groups

To dissociate the contributions of visual landmarks and path integration, we trained three groups of rats under different cue-availability conditions (Fig. 2A): (1) Path Integration + Visual Cue (PI+VC), in which both visual landmark and self-motion information were available; (2) Path Integration (PI), in which only self-motion information reliably specified the task reference frame; and (3) Visual Cue (VC), in which only visual landmark information reliably specified the task reference frame.

**Figure 2.**
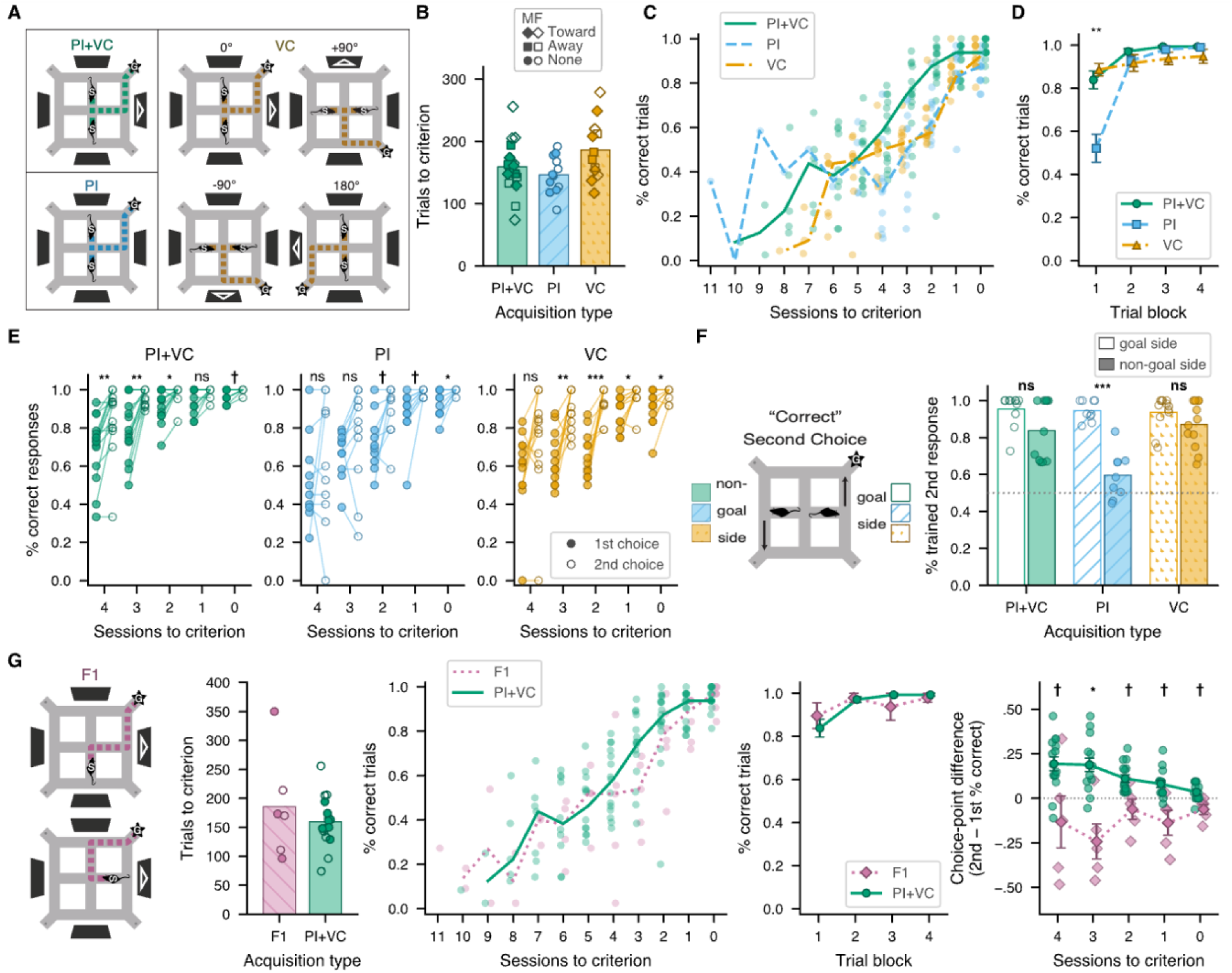
Acquisition results. (A) Training conditions. PI+VC rats were trained with both path integration and a distinguishing visual landmark available; PI rats were trained without a distinguishing visual landmark; VC rats were trained with the visual landmark while the task reference frame (visual landmark, start arms, and goal location) was pseudorandomly rotated by multiples of 90° between trials to render path integration uninformative. (B) Trials required to reach acquisition criterion. Bars show group means; symbols show individual rats, coded by sex and cue-approach direction. (C) Percentage of correct trials per session by group, aligned to the criterion session; lines show group medians, symbols show individual rats. (D) Task performance across four 8-trial blocks of the final (criterion) acquisition session; PI rats showed reduced accuracy during the first block but performed comparably to the other groups thereafter. (E) Response accuracy at the 1st (filled) vs 2nd (open) choice points over the last 5 acquisition sessions, shown per group. (F) Percentage of trained 2nd-choice responses on the goal vs non-goal sides of the maze (the trained response earns reward only on the goal side); significance markers denote within-group comparisons of responses on the goal vs. non-goal side. G) Comparison between PI+VC and F1 rats. From left: trials to criterion (as in ‘B’), percentage of correct trials per session aligned to criterion (as in ‘C’), accuracy across four 8-trial blocks of the criterion session (as in ‘D’), per-session choice-point difference score (2nd − 1st choice accuracy) with Holm-corrected PI+VC vs F1 comparisons marked above each session. †p < .10, *p < .05, **p < .01, ***p < .001; ns, not significant.

Rats in the PI+VC group (8 males, 9 females) performed the 2S2C task with a distinguishing visual landmark displayed on one of the four overhead monitors during the goal-seeking (but not the start-seeking) phase of each trial. This landmark broke the rotational symmetry of the monitor array, creating an asymmetric distal cue constellation that provided directional information and thereby specified the task reference frame. The visual landmark, goal corner, and start arms remained in fixed positions relative to one another throughout each session, allowing rats to use both visual landmarks and path integration to navigate to the goal.

Rats in the PI group (6 males, 6 females) were trained identically to the PI+VC group, with fixed start and goal locations, but no distinguishing landmark was displayed during either the goal-seeking or start-seeking phase. The four monitors therefore formed a rotationally symmetric array of distal cues that provided dim illumination without providing directional landmark information to specify the task frame. Rats consequently had to rely on path integration to determine their position and heading relative to the goal.

Rats in the VC group (7 males, 5 females) were trained with the asymmetric cue constellation present during the goal-seeking (but not the start-seeking) phase, as in the PI+VC condition. However, path integration was rendered uninformative by pseudorandom trial-wise rotations of the task reference frame (Fig. 2A, right), so that in contrast with the PI group, self-motion information accumulated during the return trajectory did not reliably predict the task configuration on the subsequent trial. This decoupling of path integration from visual cue information forced rats to rely solely on the distal landmark constellation to determine the task reference frame and select the appropriate route to the goal.

#### All groups learned at similar rates

Training sessions continued daily until rats completed 32 trials (16 from each start arm) with at least 24 correct trials. This criterion was selected to minimize overtraining while ensuring that the probability of reaching criterion by chance was ≤ .01, assuming two equiprobable response alternatives (binomial distribution, n = 32, p = .5). A correct trial was defined as one in which the rat entered the rewarded corner first, before visiting any unrewarded corner, after leaving the start arm.

Rats in the PI+VC group reached criterion after a mean of 159.4 ± 10.3 SE training trials (Fig. 2B), across a median of 7 sessions (Fig. 2C). PI rats (trained without a distinguishing visual landmark) required 146.6 ± 8.7 SE trials across a median of 5 sessions, whereas VC rats reached criterion after 186.2 ± 14.0 SE trials across a median of 7 sessions. A Kruskal–Wallis test on sessions to criterion revealed no significant effect of group (H(2) = 4.83, p = .089). A one-way ANOVA on trials to criterion similarly did not reveal a significant group effect, although there was a trend toward a difference (F(2, 38) = 2.92, p = .066), reflecting the greater number of trials required by VC rats relative to the other two groups.

#### PI rats used the goal to infer maze orientation

Fig. 2D shows task accuracy on the final training day, divided into four blocks of 8 trials each. A 4 × 3 mixed ANOVA (Greenhouse–Geisser corrected) with trial block (1–4) as a repeated factor and group (PI+VC, PI, VC) as a between-subjects factor revealed a significant main effect of group (F(2, 38) = 8.79, p < .001) and a significant block × group interaction (F(3.94, 74.8) = 12.29, p < .001). Bonferroni-corrected post hoc tests showed that this effect was driven by poorer performance in the PI group during the first trial block (vs. PI+VC, p = .007; vs. VC, p < .001); no differences involving the PI group were observed in later blocks (p > .99).

This early deficit in the PI group arose because, as noted above, the task reference frame was rotated with respect to the room before each session (to prevent rats from relying upon uncontrolled extramaze cues). Unlike the other two groups, PI rats had no asymmetric visual cue constellation to indicate the new orientation of the task frame at the start of a session; the four identical dimly backlit monitors provided no directional landmark information. PI rats therefore had to use reward feedback from early trials to identify the goal location and infer the orientation of the task reference frame. This produced reduced performance during the first trial block, but once the task-frame orientation had been inferred, PI rats performed as well as the other groups during subsequent blocks.

#### 2nd choice was learned faster than 1st choice

Fig. 2E shows mean accuracy at the 1st and 2nd choice points across the final five sessions of acquisition training for each group. Because PI rats lacked directional spatial bearings during the first trial block of each session (Fig. 2D), the first 8 trials were excluded from these calculations for all groups. Choice-point accuracy was scored independently of overall trial accuracy: after an incorrect 1st choice, rats could still reverse course before entering an unrewarded corner and thereby complete the trial correctly. Consequently, 1st-choice accuracy could fall below the acquisition criterion even when overall trial accuracy exceeded criterion.

A 2 × 5 × 3 mixed ANOVA (Greenhouse–Geisser corrected) was conducted with choice point (1st vs. 2nd) and session (1–5) as repeated factors and group (PI+VC, PI, VC) as a between-subjects factor. Only rats requiring five or more sessions to reach criterion were included (PI+VC: n = 15; PI: n = 10; VC: n = 12), because rats acquiring the task more rapidly lacked data for all five sessions. The analysis revealed a strong main effect of session (F(1.92, 65.2) = 43.4, p < .001), reflecting improved accuracy across training, and a main effect of choice point (F(1, 34) = 84.2, p < .001), reflecting greater accuracy at the 2nd than the 1st choice point. There was also a significant main effect of group (F(2, 34) = 16.4, p < .001), reflecting higher overall accuracy in PI+VC rats than in PI and VC rats (Tukey HSD, both p < .001), which did not differ from one another (p = .49). A significant group × choice point interaction (F(2, 34) = 4.29, p = .022) indicated that the magnitude of the 2nd-choice advantage differed across groups. However, planned contrasts confirmed that 2nd-choice accuracy was significantly higher than 1st-choice accuracy in all three groups (PI+VC: t(14) = 6.68, p < .001; PI: t(9) = 4.03, p = .003; VC: t(11) = 5.78, p < .001). A significant session × choice point interaction (F(2.47, 83.9) = 4.31, p = .011) indicated that accuracy gains across sessions were greater at the 1st than the 2nd choice point, consistent with the lower initial accuracy at the 1st choice point allowing greater room for improvement across sessions. No other interactions were significant (group × session F(3.84, 65.2) = 2.06, p = .10; group × choice point × session F(4.93, 83.9) = 2.21, p = .062).

#### Faster learning at 2nd choice point was not driven by proximity to the goal

One possible explanation for the faster acquisition of the 2nd choice response is that it occurred closer to the goal. In temporal-difference reinforcement learning, reward information propagates backward through preceding states and actions during learning (Sutton & Barto, 2018), which can produce faster acquisition of responses occurring closer in time or space to reward than those occurring earlier in a behavioral sequence. Alternatively, the 2nd choice may have been learned faster simply because it required the same response on every trial, whereas the correct 1st response varied with starting location.

To distinguish these possibilities, we trained a separate cohort of rats (denoted the F1 group) on the PI+VC protocol in which the correct turn at the 1st choice point remained constant across trials, while the correct turn at the 2nd varied with start location, exactly the reverse of the standard PI+VC condition (Fig. 2G). To compare PI+VC and F1 rats, we calculated a choice-point difference score (2nd-choice accuracy − 1st-choice accuracy) for each session and performed a 2 × 5 mixed ANOVA (Greenhouse–Geisser corrected) on these scores, with group (PI+VC vs. F1) as a between-subjects factor and session (1–5) as a repeated factor (Fig. 2G, right). As in Fig. 2E, the analysis was restricted to rats with at least five acquisition sessions (PI+VC, n = 15; F1, n = 5: 2 males, 3 females). Unlike the analysis in Fig. 2E, all trials from each session were included because no PI rats were included in this comparison.

The analysis revealed a significant main effect of group (F(1, 18) = 39.0, p < .001), reflecting opposite choice-point advantages in the two conditions: PI+VC rats, for which the 2nd response was fixed, showed greater accuracy at the 2nd than the 1st choice point, whereas F1 rats, for which the 1st response was fixed, showed the opposite pattern. A significant group × session interaction (F(2.58, 46.4) = 4.27, p = .013) indicated that the magnitude of this difference changed across training, diminishing as performance approached criterion. Thus, the choice-point advantage followed the location of the fixed response rather than proximity to the goal: whichever choice point required the same turn across trials was learned more rapidly. This finding is consistent with preferential acquisition of a repeated response when a stable action can be associated with a particular task state.

#### Path integration reduced perseveration of frequently rewarded responses

The 2nd choice occurred in one of two possible decision states, depending on whether the preceding 1st choice had been correct or incorrect. Following a correct 1st choice, rats encountered the 2nd choice point on the goal side of the maze, where the trained response led toward reward. Following an incorrect 1st choice, rats encountered the corresponding choice point on the non-goal side, where neither response led directly to reward.

To assess whether rats distinguished between these decision states, we analyzed the percentage of “trained” responses at the 2nd choice point on the goal versus non-goal sides of the maze (Fig. 2F). The trained response was defined as the turning response that led toward reward on the goal side and was therefore repeatedly reinforced during acquisition; the opposite turn was designated the untrained response. Values were averaged across the final five acquisition sessions for each rat and then across rats. As in Fig. 2E, the first 8 trials of each session were excluded for all groups because this block preceded spatial reorientation in PI rats. We also excluded sessions with insufficient sampling of the non-goal side (fewer than 3 incorrect 1st choices) or poor task performance (<70% accuracy at either choice point on the goal side), leaving PI+VC n = 11, PI n = 9, and VC n = 12.

A 2 × 3 mixed ANOVA with side (goal vs non-goal) as a repeated factor and group (PI+VC, PI, VC) as a between-subjects factor revealed a significant main effect of side (F(1, 29) = 43.15, p < .001), indicating that rats performed the trained 2nd response more frequently on the goal side of the maze (where it was rewarded) than the non-goal side (where it was not rewarded). A significant side × group interaction (F(2, 29) = 9.9, p < .001) further indicated that sensitivity to decision state differed across training groups. On the non-goal side, PI+VC rats performed the trained 2nd response on 83.8% of trials and VC rats on 87.1% of trials, both significantly above chance (50%; both p < .001), despite the fact that this response did not lead to reward on the non-goal side. Thus, rats trained with the visual cue constellation showed a strong tendency to perseverate on the trained 2^nd^ response even when that response did not lead toward the goal because the rat was on the wrong side of the maze.

By contrast, PI rats performed the trained 2nd response on only 59.6% of non-goal-side trials, which did not significantly exceed chance (p = .052). Thus, rats trained to rely exclusively on path integration were more sensitive to the task state in which the 2nd choice occurred and showed less perseveration on the most frequently reinforced response than rats trained with visual landmark information. This result is consistent with greater ability to discriminate between task states (visually similar T-intersections on opposite sides of the maze) in the PI group.

### Novel Route Probe Test

To assess whether rats had acquired flexible versus rigid navigation strategies, we administered a Novel Route probe session one day after acquisition criterion was reached. This probe tested whether rats could infer an optimal route to the goal from a previously untrained starting location (Fig. 3A). The session consisted of two phases. Phase 1 contained 16 familiar-route trials (8 from each of the proximal and distal trained start arms), identical to those given during acquisition. Phase 2 contained 16 Novel Route trials interleaved with 8 familiar-route trials (4 from each trained start arm). Novel Route trials therefore required rats to generalize their prior learning to a previously untrained trajectory involving a new sequence of turns. Video S2 shows a familiar-route trial and the first and last Novel Route trials of one PI_PI_ and one VC_VC_ rat.

**Figure 3.**
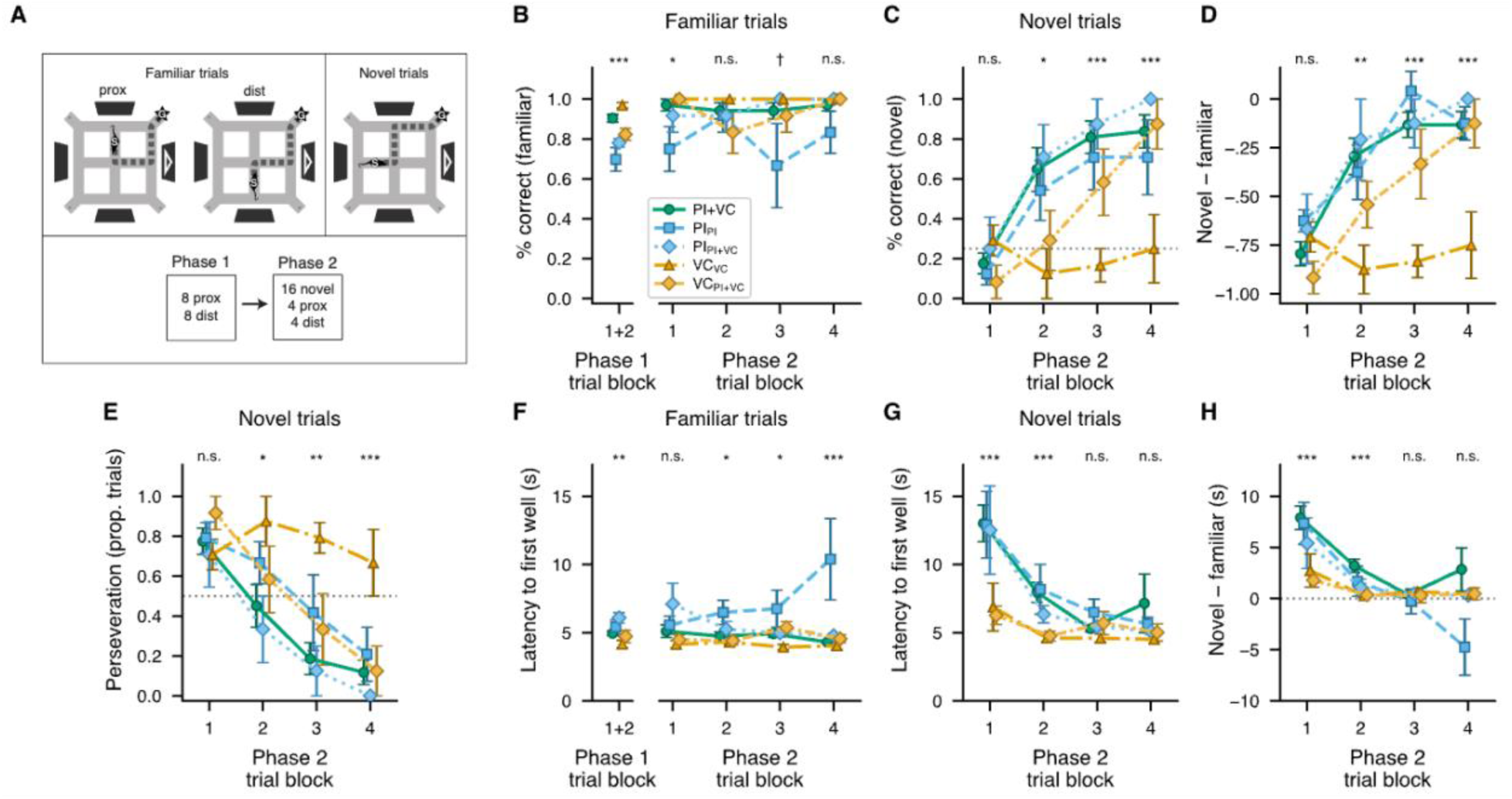
Novel Route probe test. (A) Maze diagrams illustrating familiar routes from the proximal (prox) and distal (dist) trained start arms (left) and the Novel Route configuration from a previously untrained start arm (right). The probe session consisted of Phase 1 (16 familiar-route trials: 8 prox, 8 dist), followed by Phase 2 (16 Novel Route trials interleaved with 8 familiar-route trials: 4 prox, 4 dist). (B) Mean percentages of correct familiar-route trials during Phase 1 (left) and across four trial blocks of Phase 2 (right, 2 trials per block). (C) Mean percentages of correct Novel Route trials across Phase 2 (4 trials per block). (D) Mean difference between Novel Route and familiar-route accuracy (Novel % − Familiar %) within each Phase 2 trial block (4 trials per block). Negative values indicate poorer performance on novel than familiar routes; values approaching zero indicate increasingly similar performance on the two route types. (E) Mean percentages of Novel Route trials during which rats persevered on one of the two familiar routes. (F) Mean latency of familiar-route trials during phase 1 (left) and across four trial blocks of Phase 2 (right, 2 trials per block). (G) Mean latency to reach the first well on Novel Route trials across Phase 2 (4 trials per block). (H) Mean difference between Novel Route and familiar-route latency within each Phase 2 trial block (4 trials per block). Positive values indicate longer latency on novel than familiar routes. Error bars represent SEM. Marks above each trial block give the result of the test at that block: (B, F) one-way ANOVA across the five groups; (C, D, E) contrast of VCVC vs. the other four groups (pooled-error F(1, 36)); (G, H) Welch t test comparing PI-trained (PI+VC, PIPI, PIPI+VC) with VC-trained (VCVC, VCPI+VC) rats. n.s. p ≥ .10, †p < .10, *p < .05, **p < .01, ***p < .001.

Each Novel Route trial began from a previously unused plus-maze arm that had never served as a start location during training. When the asymmetric cue constellation was present, the novel start arm was selected such that the distinguishing landmark appeared directly in front of the rat at the start of the trial (a configuration never encountered during acquisition) while the landmark’s position relative to the goal remained unchanged (rightmost maze diagram in Fig. 3A). The optimal route from the novel start to the goal therefore required a sequence of turning responses distinct from either of the two previously trained routes. Critically, successful Novel Route performance always required the response at the 2nd choice point to be opposite from the response that had previously been correct on every familiar-route trial, providing a direct test of their ability to override the previously reinforced 2nd-choice response.

#### Experimental groups

To dissociate the contributions of path integration and visual landmark information to Novel Route performance, five experimental groups were tested. The PI+VC group (n = 17) remained intact from acquisition. The PI and VC groups were each divided into two subgroups, with subscripts indicating the cue condition experienced during the Novel Route probe. PI rats (n = 12), which had been trained without a distinguishing visual landmark, were divided such that half remained in the PI condition during the probe (PI_PI_, n = 6), whereas the other half received the asymmetric visual cue constellation for the first time during the probe (PI_PI+VC_, n = 6). VC rats (n = 12), which had been trained with trial-wise rotations of the task frame that rendered path integration uninformative, were divided such that half remained in the VC condition during the probe (VC_VC_, n = 6), whereas the other half experienced a fixed task frame that rendered path integration informative for the first time while retaining the familiar visual landmark (VC_PI+VC_, n = 6). For the PI_PI+VC_ and VC_PI+VC_ groups, these changes in cue conditions were implemented throughout both Phase 1 and Phase 2 of the probe session. Sex was counterbalanced across subgroups.

#### Familiar Route performance

Performance on familiar-route trials during the probe session was analyzed to determine whether prior learning remained intact despite changes in cue conditions and the introduction of a novel start location. Fig. 3B shows mean familiar-route accuracy during Phase 1 (one block of 16 trials; left) and Phase 2 (four blocks of 2 trials; right).

A one-way ANOVA on Phase 1 accuracy revealed a significant between-subjects effect of group (F(4, 36) = 10.86, p < .001), driven primarily by reduced performance in the two groups originally trained without a distinguishing visual landmark (PI_PI_, 69.8%; PI_PI+VC_, 78.1%). This was expected since, as during acquisition (Fig. 2D), these rats had to use early trial outcomes to infer the orientation of the task reference frame. A planned contrast confirmed that PI_PI_ and PI_PI+VC_ rats performed significantly worse than groups tested with the asymmetric cue constellation (PI+VC, VC_VC_, and VC_PI+VC_; F(1, 36) = 28.33, p < .001). Bonferroni-corrected post hoc comparisons revealed no difference between PI_PI_ and PI_PI+VC_ (p = 1.0), indicating that introduction of the visual landmark did not produce a detectable improvement in familiar-route performance in rats previously trained without it. Among groups previously trained with the visual landmark, PI+VC did not differ significantly from VC_VC_ or VC_PI+VC_ (p ≥ .14), although VC_VC_ rats, which remained in their familiar rotating-frame condition, performed better than VC_PI+VC_ rats, which experienced a fixed task frame for the first time during the probe (p = .013).

A 4 × 5 mixed ANOVA (Greenhouse–Geisser corrected) on Phase 2 familiar-route accuracy, with trial block (1–4) as a repeated factor and group as a between-subjects factor, revealed no significant main effect of group (F(4, 36) = 2.18, p = .091), block (F(2.43, 87.5) = 1.06, p = .36), or block × group interaction (F(9.72, 87.5) = 1.52, p = .15). Thus, group differences were confined to Phase 1 and the first Phase 2 block (Fig. 3B); thereafter, familiar-route performance was comparable across all groups and remained stable during Phase 2 despite the interleaved Novel Route trials.

#### Novel Route performance

If rats had acquired a map-based representation of the goal location, then they might be able to immediately infer an appropriate route from a previously untrained starting location. By contrast, rats relying on a rigid route-based strategy might initially repeat previously reinforced response sequences even though those responses do not lead to the goal from the novel start location.

Fig. 3C plots mean percentages of correct Novel Route trials across the four blocks of Phase 2. Accuracy during the first block was low in every group (PI+VC: 17.6 ± 5.1%; PI_PI_: 12.5 ± 5.6%; PI_PI+VC_: 25.0 ± 15.8%; VC_VC_: 29.2 ± 7.7%; VC_PI+VC_: 8.3 ± 8.3%), indicating that rats did not immediately infer the optimal route from the novel start arm. To assess each rat’s Novel Route performance relative to its own baseline performance on familiar routes, Fig. 3D plots the difference between each rat’s percentage of correct Novel and familiar route trials (Novel − Familiar) within each block of Phase 2. Negative values indicate poorer performance on novel than familiar trials. A 4 × 5 mixed ANOVA revealed a significant main effect of block (F(3, 108) = 30.69, p < .001), indicating that Novel Route performance improved across Phase 2 and became increasingly similar to familiar-route performance. A main effect of group (F(4, 36) = 6.50, p < .001) combined with a significant block × group interaction (F(12, 108) = 3.15, p < .001) indicated that improvement differed across groups. Groups with prior experience using path integration approached asymptotic Novel Route performance earlier in the session, whereas VC_PI+VC_ rats, for which path integration became informative only during the probe, improved more slowly. VC_VC_ rats, which were unable to use path integration during training or the probe test, were the only group that showed no improvement in Novel Route performance across the entire session.

#### Perseveration of Familiar Responses

All groups had a high error rate on early Novel Route trials. To determine whether these errors reflected perseveration of previously learned responses, we scored each Novel Route trial according to whether the rat repeated one of the two response sequences reinforced during acquisition (Fig. 3E), neither of which led directly to the goal from the novel start arm. A small number of trials for which the route could not be classified were excluded (24/656 trials, 3.7%).

Regardless of training condition, rats initially tended to fall back on previously reinforced response sequences when first confronted with the Novel Route. During the first block, mean perseveration exceeded 70% in every group (PI+VC: 50/62, 81%, p < .001; PI_PI_: 18/23, 78%, p = .011; PI_PI+VC_: 17/23, 74%, p = .035; VC_PI+VC_: 21/23, 91%, p < .001; VC_VC_: 17/24, 71%, p = .064), with no significant difference among groups (F(4, 36) = 0.62, p = .65).

Groups differed markedly, however, in how readily they abandoned these responses as the session progressed. A 4 × 5 mixed ANOVA on per-rat perseveration revealed significant main effects of block (F(3, 108) = 33.56, p < .001) and group (F(4, 36) = 3.43, p = .018), as well as a block × group interaction (F(12, 108) = 2.19, p = .017). From block 1 to block 4, perseveration declined from 81% to 12% in PI+VC, 78% to 23% in PI_PI_, 74% to 0% in PI_PI+VC_, and 91% to 12% in VC_PI+VC_, but remained high in VC_VC_ (71% to 67%). A contrast comparing VC_VC_ with all other groups across the session supported this dissociation (F(1, 36) = 9.28, p = .004; Welch t(6.9) = 3.21, p = .015). Thus, groups differed not in their initial tendency to perseverate, but in their subsequent ability to abandon previously reinforced response sequences. Only VC_VC_ rats, which lacked access to informative path-integration signals during the probe, continued to perseverate throughout the entire session.

#### Latency to reach the first well

We next measured the latency from trial initiation (start-arm door opening) until arrival at the first corner well, regardless of whether it was the rewarded goal. Fig. 3F shows familiar-route latencies during Phases 1 and 2. Mean Phase 1 latencies were approximately 5 s in all groups, although PI-trained groups tended to be slightly slower than VC-trained groups, producing a significant group effect (F(4, 36) = 4.15, p = .007). Familiar-route latencies remained relatively stable during Phase 2 in most groups, although several PI_PI_ rats exhibited large increases during later blocks, producing substantial heterogeneity; these differences were not pursued further.

Despite similarly poor initial Novel Route accuracy and similarly high perseveration across groups (Figs. 3C–E), Novel Route latencies revealed a striking difference between PI- and VC-trained rats (Figs. 3G,H). During the first block, VC-trained rats reached the first well in about 6.5 s (median 5.4 s), roughly 2 s slower than on familiar-route trials, whereas PI-trained rats required nearly 13 s, more than twice their familiar-route latencies (Fig. 3G; Video S2). To quantify this slowing relative to familiar-route performance, Fig. 3H plots the difference between each rat’s Novel and familiar-route latencies (Novel − Familiar) within each Phase 2 block. Positive values indicate longer latencies on novel trials. A 4 × 5 mixed ANOVA (Greenhouse–Geisser corrected) revealed significant effects of block (F(1.90, 68.4) = 12.46, p < .001) and group (F(4, 36) = 2.65, p = .049), but no significant block × group interaction (F(7.60, 68.4) = 1.72, p = .11). Planned Welch comparisons between pooled PI-trained groups (PI+VC, PI_PI_, PI_PI+VC_) and VC-trained (VC_PI+VC_, VC_VC_) rats showed greater Novel Route slowing in PI-trained rats during blocks 1 (PI: 7.28 s, VC: 2.26 s; t(33.8) = 3.99, p < .001) and 2 (PI: 2.46 s, VC: 0.32 s; t(30.1) = 4.02, p < .001), but not blocks 3 (PI: 0.26 s, VC: 0.49 s; t(27.1) = −0.47, p = .64) or 4 (PI: 0.75 s, VC: 0.46 s; t(30.4) = 0.19, p = .85).

In summary, although PI- and VC-trained rats initially showed similarly poor Novel Route accuracy and high perseveration, their response to the novel starting condition differed markedly. PI-trained rats exhibited pronounced slowing of their behavior during early Novel Route trials that disappeared as Novel Route performance improved, whereas VC-trained rats responded rapidly on all Novel Route trials. This pattern suggests that PI-trained rats responded to the novel situation of the new start arm with increased deliberation before adapting their behavior, whereas VC-trained rats continued to execute learned responses habitually with little deliberation.

### Reversal Probe Test

To further evaluate the flexibility of rats’ navigation strategies, we administered a Reversal probe session following the Novel Route probe. This session tested whether rats could adapt to a relocated goal while navigating from the two originally trained start arms (Fig. 4A; Video S3). Phase 1 consisted of 16 familiar-route trials (8 from each start arm) with the goal in its original location. In Phase 2, the goal was relocated to the opposite corner for up to 64 Reversal trials (32 from each start arm). The required turning sequences themselves remained unchanged: one route required repeated turns and the other alternating turns. However, their mapping to the two start arms was reversed, such that the start arm that previously required a repeated response sequence now required an alternating sequence, and vice versa.

**Figure 4.**
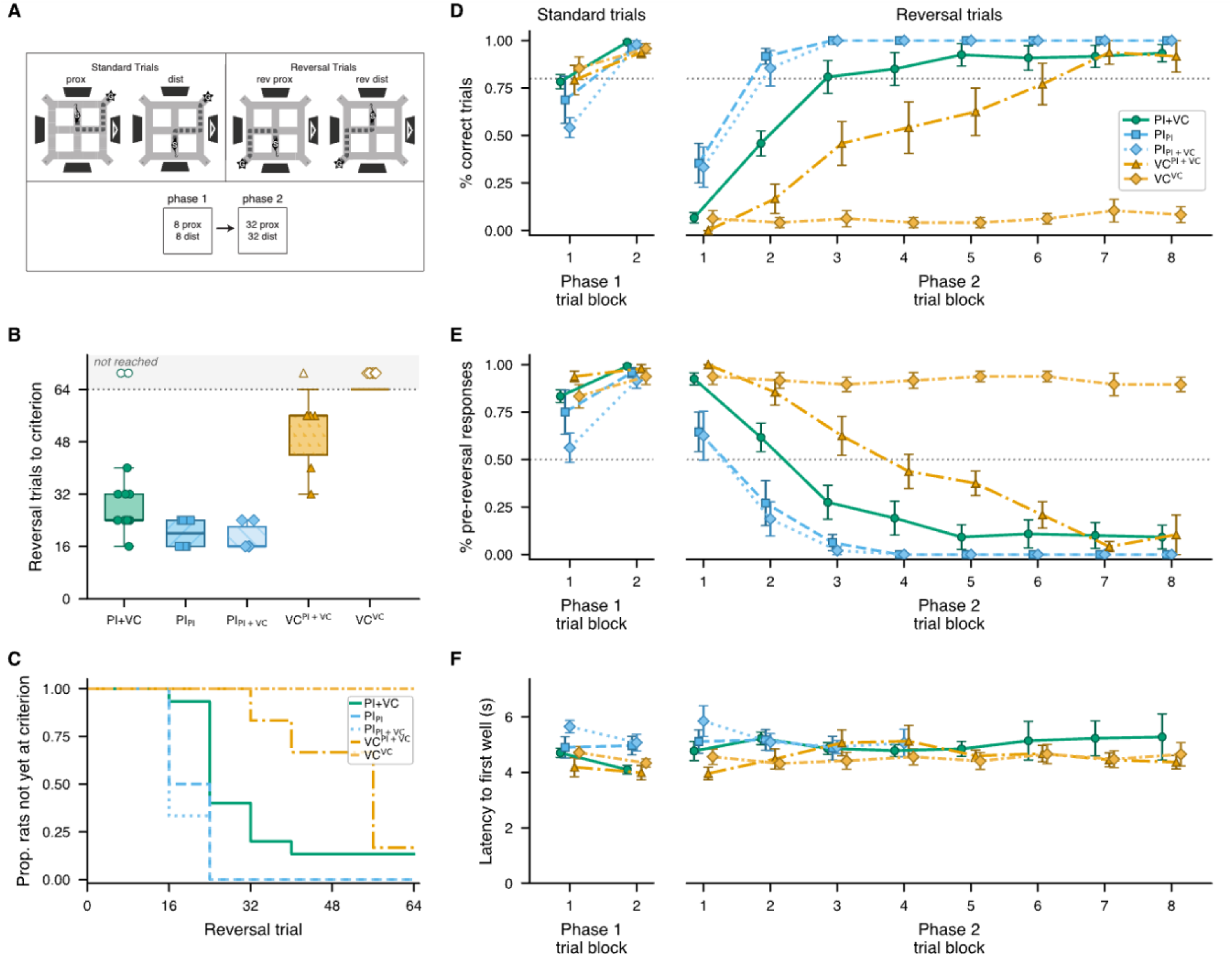
Reversal probe test. (A) Maze diagrams illustrating familiar routes from the proximal (prox) and distal (dist) trained start arms (left) and Reversal routes after the goal was relocated to the opposite corner (right). Repeated and alternating turning sequences remained the same as during acquisition, but their mapping to the two start arms was reversed. Phase 1 consisted of 16 familiar-route trials (8 prox, 8 dist) with the goal in its originally trained location; Phase 2 consisted of up to 64 Reversal trials (32 rev prox, 32 rev dist). (B) Box-and-whisker plots (median, interquartile range, whiskers to 1.5 × IQR) of the number of reversal trials each rat needed to reach a criterion of 8 correct trials within an 8-trial block; symbols show individual rats. Open symbols in the shaded band mark rats that had not reached criterion by the end of the 64 reversal trials. (C) Percentage of rats that had not reached a criterion of 8 correct trials within an 8-trial block by the end of each trial block. (D) Mean percentage of correct trials across Phase 1 (two blocks of 8 trials) and Phase 2 (eight blocks of 8 trials) for the five experimental groups. VC subgroups are identified by superscripts indicating their task-frame condition during Reversal (VCPI+VC, stable frame; VCVC, rotating frame). Both PI subgroups underwent Reversal without a distinguishing visual landmark and are identified by subscripts indicating their differing prior Novel Route conditions (PIPI+VC, visual landmark during Novel Route; PIPI, no visual landmark during Novel Route), allowing influences of prior cue exposure to be tracked despite identical Reversal conditions. (E) Block-wise percentage of trials on which rats made the pre-reversal (familiar) first turn; as in D, rats that stopped after reaching criterion were scored as 0% for their remaining blocks. (F) Block-wise mean latency from trial start to the first well entry; group × block cells with fewer than three rats are not shown. Error bars represent SEM.

#### Experimental groups

Five groups were tested during the Reversal probe to dissociate the contributions of path integration and visual landmark information to reversal learning. The PI+VC group (n = 15; two further PI+VC rats were excluded from the Reversal analyses, see Reversal performance) was carried forward from acquisition and the Novel Route probe (these rats also completed two intervening probe sessions; see Methods and Results). VC-trained rats were divided according to the task-frame condition that they had previously experienced during the Novel Route Probe. VC^PI+VC^ rats (n = 6, formerly VC_VC_ rats which had remained in the rotating task frame during acquisition and the Novel Route probe), experienced a stable task frame during Reversal, making path integration informative for the first time during the Reversal session. Conversely, VC^VC^ rats (n = 6, formerly VC_PI+VC_ rats which had experienced a stable task frame for the first time during the Novel Route probe), returned to the rotating-frame condition during Reversal, rendering path integration uninformative once again. All PI-trained rats were tested without a distinguishing visual landmark during Reversal (the same condition under which they had originally been trained), but since these rats had been subdivided into two groups (PI_PI+VC_ and PI_PI_) during Novel Route probes, they were analyzed as separate subgroups to track whether their differing experience during the prior probe session affected their subsequent behavior during identical reversal probes.

#### Pre-reversal performance

Fig. 4D (left) shows mean trial accuracy during Phase 1 (two blocks of 8 trials each), when the goal remained in its originally trained location. A 2 × 5 mixed ANOVA with block (1 vs. 2) as a within-subjects factor and group (PI+VC, PI_PI+VC_, PI_PI_, VC^PI+VC^, VC^VC^) as a between-subjects factor revealed a modest block × group interaction (F(4, 34) = 2.98, p = .033). Bonferroni-corrected paired comparisons (five tests) showed that between blocks 1 and 2, PI_PI+VC_ rats improved from 54% to 98% (t(5) = 7.00, p = .005) and PI+VC rats from 78% to 99% (t(14) = 5.23, p < .001), whereas the improvement in PI_PI_ rats (69% to 96%) did not reach significance (t(5) = 2.14, corrected p = .43), and both VC subgroups, which were already at 79–85% in block 1, showed no significant change (corrected p ≥ .42). The improvement observed in PI-trained groups is again consistent with these rats using early trial outcomes to re-infer the orientation of the task reference frame from the goal location, as observed during acquisition and the Novel Route probe (Figs. 2D, 3B).

#### Reversal performance

Fig. 4B shows that groups differed significantly in the median number of trials needed to reach a criterion of 8 correct trials within a block of 8 reversal trials (Kruskal-Wallis H(4) = 26.98, p < .001; nine rats never reached criterion within the 64 reversal trials: all six VC^VC^ rats, two PI+VC rats and one VC^PI+VC^ rat). This criterion was selected such that the probability of reaching criterion by chance was ≤ .01, assuming two equiprobable response alternatives (binomial distribution, n=8, p=0.5). Fig. 4C shows the percentage of rats in each group that had failed to reach criterion by the end of each trial block in Phase 2 (eight blocks of 8 trials each).

Fig. 4D shows mean trial accuracy during each block of Phase 2. Some rats were removed from the maze after reaching criterion, and for these rats, a reversal performance of 100% was assumed in each block after criterion had been reached. Two PI+VC rats whose sessions were ended before they had met this criterion could not be scored in this way and were excluded from all Reversal analyses (final n = 15 for PI+VC). An 8 × 5 mixed ANOVA on Phase 2 accuracy (Greenhouse–Geisser corrected) revealed a highly significant main effect of block (F(2.75, 93.4) = 84.9, p < .001), reflecting successful reversal learning in most groups. However, a significant main effect of group (F(4, 34) = 32.7, p < .001) and block × group interaction (F(10.99, 93.4) = 8.74, p < .001) indicated substantial group differences in the rate and extent of reversal learning.

PI-trained rats reversed most rapidly. Both PI_PI+VC_ and PI_PI_ groups exceeded 80% mean accuracy by the second trial block and reached criterion in fewer trials than PI+VC rats (medians 16 and 20 vs. 24 trials; Holm-corrected Mann–Whitney tests, p = .042 and .034) and showed higher block-averaged Phase 2 accuracy, significantly so for PI_PI_ (PI_PI_ vs. PI+VC, Bonferroni-corrected p = .046; PI_PI+VC_ vs. PI+VC, p = .081). Prior exposure to the visual landmark during the Novel Route probe therefore did not prevent rapid reversal when rats were subsequently tested under PI-only conditions. PI+VC rats reversed more gradually, requiring three trial blocks (24 trials) to exceed 80% mean accuracy.

VC-trained rats showed the poorest reversal performance. VC^VC^ rats, which were trained and reversed with a rotating task frame that rendered path integration uninformative, failed to reverse and instead continued to perseverate on pre-reversal responses throughout all eight blocks (Fig. 4E; Video S3). Their mean Phase 2 accuracy (6%) was lower than that of every other group (independent-samples t tests, Welch-corrected for unequal group sizes; all Bonferroni-corrected p < .002). By contrast, VC^PI+VC^ rats, which had also been trained with a rotating task frame but experienced a stable frame during Reversal, eventually acquired the reversal (five of six rats reached criterion; group accuracy first exceeded 80% in block 7), but more slowly than the other groups that reversed (median 56 trials to criterion vs. 24 for PI+VC and 16–20 for the PI subgroups; Holm-corrected Mann–Whitney tests, all p ≤ .042). Fig. 4F shows that mean trial latencies remained within the range of 4–6 seconds throughout the Reversal session for all groups.

In summary, reversal performance depended strongly on both training history and the information available during the Reversal probe. Rats trained to rely on path integration reversed rapidly. Rats trained to rely exclusively on visual landmarks failed to reverse when path integration remained uninformative in Reversal, but making path integration informative for the first time during Reversal rescued learning in VC-trained rats, albeit only gradually.

### Individual Differences in Navigational Strategy

As noted above, rats trained in the PI+VC condition (n = 17) completed two additional probe sessions after the Novel Route probe and before the Reversal probe: a No-Cue probe, equivalent to the PI acquisition condition, and a Rotation probe, equivalent to the VC acquisition condition. To characterize individual differences in navigational strategy, we represented each rat in a three-dimensional performance space defined by accuracy on the Novel Route, No-Cue, and Rotation probes (Fig. 5A). Reversal performance was withheld from this analysis for use as an independent external validation measure.

**Figure 5.**
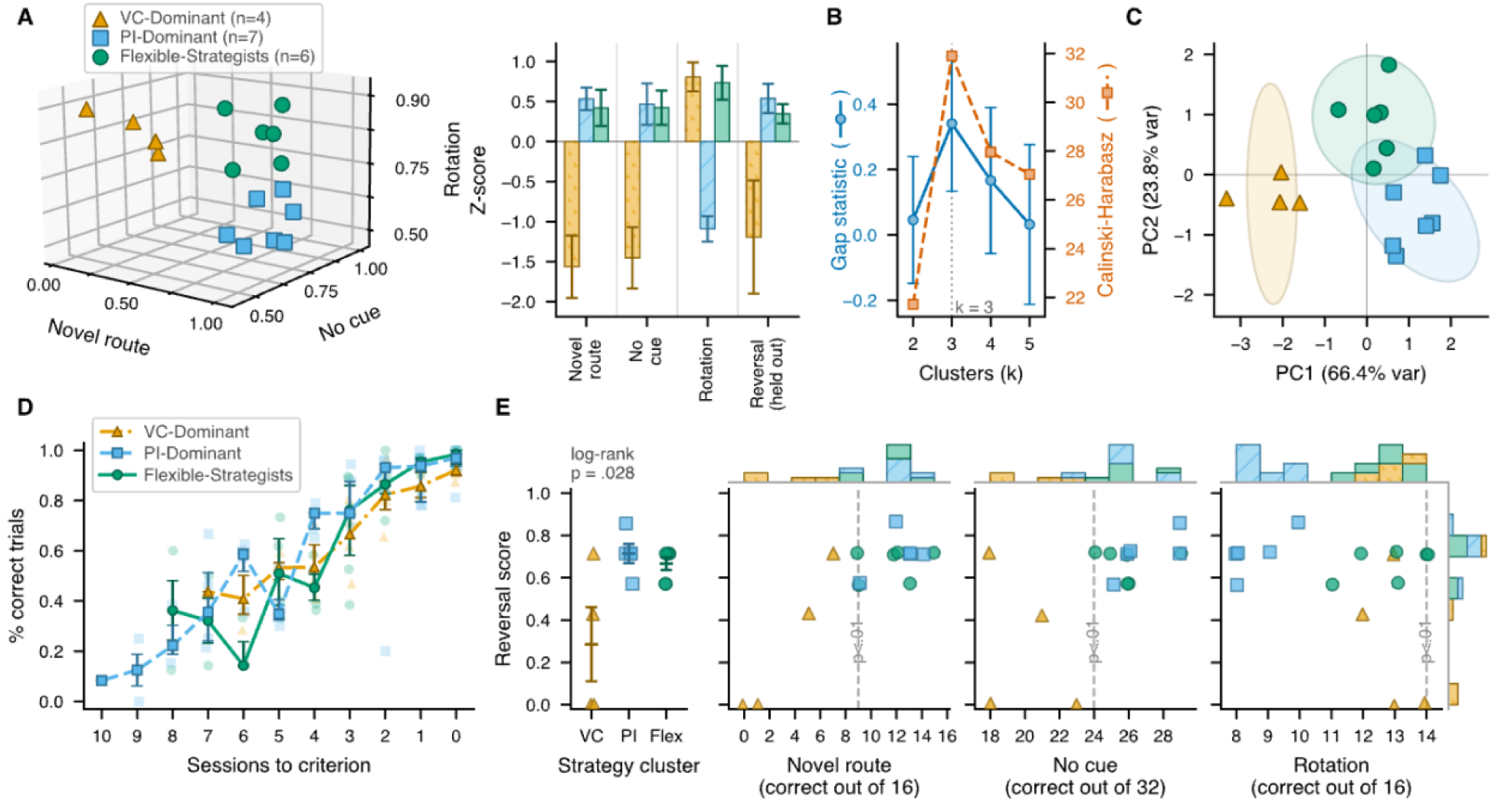
Individual differences in navigational strategy among PI+VC rats. (A) Left: 3D plot of each rat’s % correct trials across the Novel Route, No-Cue, and Rotation probes, coded by behavioral strategy profile identified by k-means clustering (k = 3). Right: Z-score normalized performance (mean ± SEM) on probe tests for each strategy cluster (Reversal probe was held out from cluster analysis and is shown for the 15 rats scored; see main text). VC-Dominant rats show low scores on path-integration measures (Novel Route, No Cue) and poor Reversal performance, but high scores on cue-following (Rotation probe). PI-Dominant rats show the opposite pattern. (B) The gap statistic (left axis; 500 reference datasets) and Calinski-Harabasz index (right axis) both peak at k = 3 strategy clusters from the probe test data. (C) PCA was performed on the three clustering variables (Novel Route, No Cue, Rotation). PC1 clearly separates VC-Dominant rats from the other two clusters; PC2 distinguishes Flexible-Strategists from PI-Dominant rats. Shaded regions denote 95% confidence ellipses. (D) Acquisition learning curves aligned to the criterion session (session 0). Lines show median percentage correct for each session, error bars show interquartile ranges. (E) Left: Reversal score by strategy cluster (mean ± SEM; p from a log-rank test on trials to criterion with never-reversed rats censored). Right: Reversal score (y-axis; trials to criterion rescaled so that 1 = criterion in the first block of 8 and 0 = not reached within 64 trials; withheld from the clustering analysis as an independent measure of behavioral flexibility; n = 15, see main text) is plotted against performance on each of the three probe tests used for clustering (Novel Route, No Cue, Rotation). Vertical dashed lines mark above-chance performance on each probe (one-sided binomial test, p < .01; chance = .25 for Novel Route, .5 for No Cue and Rotation).

#### Three behavioral strategy profiles

K-means clustering was applied to the first 2 principle components of the three-dimensional performance data. A three-cluster solution was adopted because both the Gap statistic (500 reference datasets) and the Calinski–Harabasz index were maximal at k = 3 (Fig. 5B), yielding three separable regions of performance space (Fig. 5C). The clustering solution was highly stable across several measures: (1) mean bootstrap assignment stability was 95.9% across 1,000 resamples (k-means refit in the principal-component space, all 17 rats reassigned); (2) cluster-wise Jaccard similarity coefficients were 0.982, 0.906, and 0.920, all exceeding the 0.75 criterion for stable clusters (Hennig, 2007); (3) leave-one-out LDA classification accuracy was 100% (17/17); and (4) removal of any individual rat did not alter the cluster assignments of the remaining animals. Cluster membership was unrelated to sex or to cue-approach direction (both χ²(2) = 1.90, p = .39; the two contingency tables coincide).

Based on results of the cluster analysis, we refer to the three behavioral strategy profiles (Fig. 5A): 1) the *VC-Dominant* profile (n = 4) was characterized by strong Rotation-probe performance (mean = 13.0/16 correct) but poor Novel Route performance (mean = 3.25/16) and lower No-Cue performance (mean = 20.0/32), consistent with greater reliance on visual landmark information than on path integration; 2) the *PI-Dominant* profile (n = 7) showed strong Novel Route (mean = 12.1/16) and No-Cue performance (mean = 26.1/32), together with lower Rotation performance (mean = 8.7/16), consistent with greater reliance on path integration than on visual landmark information; 3) The *Flexible-Strategist* profile (n = 6) performed strongly across all three probes (Novel Route: mean = 11.7/16; No Cue: mean = 26.0/32; Rotation: mean = 12.8/16), consistent with an ability to exploit either path-integration or visual-landmark information according to task demands. Rats with different strategy profiles acquired the task at similar rates during training (Fig. 5D; sessions to criterion, Kruskal–Wallis H(2) = 0.08, p = .96; trials to criterion, H(2) = 2.48, p = .29).

#### Relationship to reversal performance

Reversal performance was withheld from the clustering analysis to provide an independent measure of behavioral flexibility (Fig. 5E). Reversal performance was scored as trials to criterion (the criterion of 8 correct trials within a block of 8 defined above), rescaled so that 1 denotes criterion in the first block and 0 denotes failure to reach criterion within 64 trials [(64 − trials to criterion)/56]; the two PI+VC rats excluded from the Reversal analyses were excluded here as well (n = 15). The resulting pattern was consistent with the strategy profiles: VC-Dominant rats showed substantially lower mean Reversal performance (0.29) than either Flexible-Strategist (0.67) or PI-Dominant rats (0.71). This difference should be interpreted descriptively, however, because the VC-Dominant cluster was small (n = 4) and heterogeneous: two rats failed Reversal entirely, whereas the other two achieved moderate performance (0.43 and 0.71). Consequently, within-cluster variability was substantially greater for VC-Dominant rats (SD = 0.35) than for the Flexible-Strategist (SD = 0.07) or PI-Dominant (SD = 0.10) groups, violating the equal-variance assumption (Levene’s test, p = .005). Although a conventional one-way ANOVA was significant (F(2, 12) = 6.64, p = .012, η² = .53), neither Welch’s ANOVA (F(2, 5.8) = 2.59, p = .16) nor the Kruskal–Wallis test (H(2) = 5.30, p = .071) reached significance. A log-rank test on trials to criterion, which treats the two rats that never reached criterion as censored rather than as scoring zero, did indicate a cluster difference (χ²(2) = 7.14, p = .028; VC-Dominant vs the other two clusters, χ²(1) = 6.66, p = .010). We therefore regard the lower Reversal performance of VC-Dominant rats as supported, although the small size and heterogeneity of this cluster limit the precision of the estimate.

Across individual rats, Novel Route performance was the strongest predictor of Reversal performance in a multiple regression model (R² = .76; HC3 heteroscedasticity-robust standard errors: Novel Route b = 0.84, t(11) = 2.47, p = .031; No Cue b = −0.19, p = .85; Rotation b = 0.04, p = .88), and the rank-order association was also significant (Spearman ρ = .63, p = .013). This association suggests that the ability to navigate from a novel starting location and the ability to adapt to a novel goal location may depend on common processes supporting flexible spatial generalization. However, because this relationship depended on the two VC-Dominant rats that never reached criterion (without them, R² = .33 and the Novel Route slope was not significant, p = .23), this interpretation should also be regarded as provisional. Taken together, these analyses suggest that even among rats receiving identical PI+VC training, individual differences in reliance on visual landmarks versus path integration were associated with differences in navigational flexibility.

## DISCUSSION

Here, we found that the source of spatial information rats relied upon during maze learning strongly influenced the flexibility of their subsequent navigational decisions. When rats were able to rely on path integration, they adapted readily to novel routes and altered goal contingencies, and they rapidly suppressed previously reinforced responses that no longer led to reward. By contrast, despite showing similar acquisition of the trained task, rats forced to rely on visual landmarks showed greater perseveration and more rigid response retrieval when challenged to follow novel routes or adapt to relocated goals. These findings suggest that reliance upon path integration can promote the acquisition and use of mental representations that capture latent spatial structure, thereby biasing navigational decisions toward flexible inference rather than rigid response repetition.

### State inference during spatial navigation

When making a navigational decision, animals must represent their current task state in a way that provides sufficient information for selecting an appropriate locomotor action. Spatial variables such as allocentric position and direction can be highly informative components of such a state representation, especially in novel situations where navigational choices depend flexibly upon spatial variables. However, allocentric spatial variables are inherently “latent”: they cannot be directly perceived by the senses, so their availability for guiding navigation depends upon whether prior experience has provided sufficient information to infer allocentric spatial states. Reliance upon path integration may provide such experience by imposing a particular representational demand: requiring task states to be inferred from temporal context.

#### Dependence of state upon temporal context

Task-state representations can differ in their dependence on temporal context. At one extreme, a state may be identified from current sensory observations alone (such as the presence of a familiar landmark), allowing new actions to be selected without reference to the sequence of events that preceded them. Such context-free representations are independent of history: the same observation evokes the same state representation regardless of how it was reached. Alternatively, state identity may depend partly or entirely on temporal context, such that the same current observation represents different task states depending on the sequence of preceding observations and actions. Such context-dependent representations give rise to the well-known “long-range dependency problem” encountered in many sequence learning domains (Bengio et al., 1994). For example, in the domain of language, interpreting an ambiguous word’s meaning may depend on previous words encountered earlier in a sequence (that is, the linguistic context). Similarly, when navigating via path integration, estimating current position may depend upon knowing previous positions encountered earlier along a trajectory.

Our findings suggest that path integration, which requires task states to be discriminated from temporal context rather than from current sensory observations alone, can promote the formation and use of task representations that encode latent allocentric spatial variables and thereby support flexible navigation in novel situations. This interpretation is consistent with prior theories proposing that spatial representations emerge from learning the sequential and relational structure of experiences (McNaughton et al., 1996, 2006; Whittington et al., 2022).

#### Contextual aliasing fosters context-free state representations

In the VC version of the 2S2C task, trial-wise rotation of the maze reference frame caused the same task states to be preceded by different self-motion histories. For example, at the beginning of a trial, the same start location (proximal or distal) could be preceded by different trajectories depending upon where the goal was located on the prior trial. This “contextual aliasing” (whereby the same task state was preceded by different temporal contexts) may have pressured VC rats to discriminate task states using learned associations with distal landmarks independently of movement history (Knierim & Hamilton, 2011; McNaughton et al., 1996). Such training may encourage a form of “behavioral overfitting,” in which a strategy accurately solves the trained problem but generalizes poorly when routes or goal contingencies change. Consistent with this interpretation, VC rats showed several indicators of rigid navigation: repetition of the trained second response on the non-goal side despite the absence of reward, perseveration on previously reinforced sequences during Novel Route trials, and failure to learn Reversal within a single session.

#### Perceptual aliasing fosters context-dependent state representations

In the PI version of the task, quadrilateral symmetry of the maze environment caused similar sensory observations to occur in functionally distinct task states. For example, the first choice points on the two start arms were observationally identical, as were the second choice points on the goal and non-goal sides of the maze. This “perceptual aliasing” (whereby the same sensory observations were experienced in different task states) may have pressured rats to discriminate task states using their memory of preceding trajectories, rather than ambiguous visual landmarks. Accordingly, several indicators of flexible navigation were observed in PI rats: minimal perseveration on inappropriate responses, rapid acquisition of Novel Routes, and fast adaptation to a new goal location during Reversal. This interpretation is consistent with findings from dual-strategy plus-maze tasks, in which rats typically transition from a map-based “place” strategy early in training to a route-based “response” strategy with overtraining (Ritchie et al., 1950; Hicks, 1964). This transition can be prevented despite extensive training when sequential navigational choices require spatial working memory for preceding decisions (Gardner et al., 2013). Like the PI condition in the present study, such a requirement makes the identity of the current decision state dependent upon information carried forward from preceding experience.

### Landmark guidance and navigational flexibility

In the 2S2C task, inflexible navigation in the VC condition was guided by visual landmarks. However, we do not infer from this that visually guided navigation is generally inflexible. Rather, we propose that inflexibility arises when navigation is controlled by context-independent state representations. In natural environments, visual landmarks may contribute to flexible navigation strategies that are based on context-dependent spatial representations.

The hippocampal system is widely recognized as a primary neural substrate for flexible, map-based navigation (O’Keefe & Nadel, 1978). Consistent with a role for path integration in flexible navigation, substantial evidence implicates the hippocampus and related structures in path-integration-dependent spatial behavior and representations (Whishaw et al., 1997; Golob & Taube, 1999; Jayakumar et al., 2019; but see Alyan & McNaughton, 1999). In contrast, inflexible response-based navigation depends more heavily on striatal action systems (Morris et al., 1982; Packard & McGaugh, 1992; McDonald & White, 1993).

The functional dissociation between these neural systems does not map directly onto path integration versus visual guidance. The hippocampus is critical not only for path integration, but also for navigation that relies upon configurations of distal visual landmarks that specify allocentric locations (Morris et al., 1982; Packard & McGaugh, 1992). Thus, visual landmark guidance can support flexible navigation and spatial representations associated with hippocampal function. One possible reason for this could be that learning relationships between landmark configurations and allocentric locations requires context-dependent representations of temporal changes in landmark configurations as animals move through an environment, although the present experiments do not address this possibility.

It remains unclear how VC rats used visual information in the 2S2C task. The four overhead monitors formed a spatially distributed distal array, resembling landmark configurations capable of supporting map-based navigation. However, rats may instead have treated the distinguishing landmark on one monitor as a beacon or discriminative cue associated directly with particular responses. Under this interpretation, reduced flexibility in VC rats could reflect reliance on a striatal cue–response system rather than, or in addition to, reduced temporal-context dependence of their state representations. These accounts are not necessarily mutually exclusive: training that made trajectory context unreliable may itself have favored a cue–response representation supported by striatal circuitry. Future studies manipulating the spatial configuration of visual cues and testing hippocampal versus striatal dependence of PI and VC performance could distinguish these possibilities.

### Flexible strategy switching during reversal

Reversal trials began from familiar locations within the standard maze configuration, but previously rewarded actions failed to produce the expected outcome and reward was subsequently encountered elsewhere. Flexible reversal therefore required rats to use the changed outcome to modify future decisions. As in Novel Route trials, access to path integration was associated with greater flexibility: PI rats reversed most rapidly, PI+VC rats more slowly, whereas VC rats tested without informative path integration continued to perseverate.

The rapid reversal of PI rats is consistent with the possibility that their state representations separated the latent spatial structure of the task from the particular actions and outcomes associated with it. Rather than acquiring an entirely new response policy, PI rats may have reassigned the goal within an existing cognitive map and then inferred the actions needed to reach it. This resembles a process they already performed at the beginning of each session: identifying the current goal location and navigating relative to it using path integration. PI+VC rats may have reversed more slowly because they additionally had to overcome learned associations between visual landmarks and rewarded responses. If rapid PI reversal reflects updating an existing spatial representation rather than gradual relearning of reinforced responses, it may be accompanied by rapid changes in hippocampal place and head-direction representations together with changes in prefrontal representations involved in outcome updating and behavioral control (Izquierdo, 2017; Schoenbaum et al., 2009), a prediction that is testable in future studies.

### Individual differences in navigational strategy

Individual differences within the PI+VC group (Fig. 5) demonstrate that identical training conditions need not produce identical strategies. Classic dual-solution paradigms typically classify animals as place or response learners from a single probe trial (Tolman et al., 1946; Packard & McGaugh, 1996), although the expressed strategy can depend on information and task demands at testing (Restle, 1957). Our multi-probe 2S2C task design instead characterized individual strategies across independent probe challenges.

Novel Route, No-Cue, and Rotation performance identified three profiles among identically trained PI+VC rats. VC-Dominant rats relied primarily on landmarks and generalized relatively poorly; PI-Dominant rats relied primarily on path integration and generalized flexibly; and Flexible-Strategist rats exploited either information source according to task demands. Reversal performance, withheld from clustering, was consistent with these profiles, although the small and heterogeneous VC-Dominant group warrants caution. Importantly, individual differences converged with the effects of experimentally constraining spatial information: forced landmark reliance produced a rigid profile resembling VC-Dominant rats, whereas forced path-integration reliance produced the flexible profile characteristic of PI-Dominant rats. Thus, informational constraints strongly biased the representations and strategies acquired, but did not uniquely determine them: when multiple information sources were available, individual rats appeared able to construct different solutions to the same navigational problem.

### Limitations, future directions, and conclusions

An important limitation of our study is that the present behavioral results do not directly reveal the internal representations or neural mechanisms underlying flexible navigation. Behavioral findings alone cannot determine whether rats explicitly represented allocentric locations, how state transition structure was encoded, or how such representations guided action selection. In future studies using the 2S2C task, neural recordings and manipulations could address these questions by testing contributions of hippocampal and prefrontal systems implicated in flexible spatial behavior versus striatal systems associated with response-based navigation (Ragozzino et al., 1999; Packard & Goodman, 2013; Patai & Spiers, 2021).

The distinct behavioral profiles within the PI+VC condition could be further leveraged to investigate whether different navigation strategies are associated with distinct neural signatures, a question that can become obscured when data are averaged across all individuals in the same treatment group (Flagel et al., 2011; Sarter & Phillips, 2018). Because rats with different profiles achieved equivalent mastery of the trained task despite markedly different generalization profiles, the 2S2C task may provide a useful model for investigating how similar behavioral outcomes arise from different underlying representations and computational strategies.

More broadly, our findings suggest that requiring temporal context for state inference may promote representations of latent spatial structure that support flexible navigational decisions. Informational constraints during learning may therefore determine not only how successfully an organism learns a task, but what it learns about the structure of that task and how flexibly that knowledge can be applied when circumstances change.

## MATERIALS AND METHODS

### Subjects

46 Long-Evans rats acquired from Charles River (23 males, 23 females, two to three months old). Animals were housed in a temperature and humidity controlled vivarium with a 12-12 reverse light-dark cycle and fed ad lib until they reached >300g for males and >200g for females, after which they were reduced to 85% of their ad lib weight and maintained there by controlled daily feeding. Prior to being part of the experiment rats were handled for 10-minute intervals on five consecutive days. When rats were ready for running the experiments, their weight was further reduced to 75% ad lib and maintained.

### Apparatus

Rats performed a novel two-start-two-choice (2S2C) task on a custom-built apparatus consisting of a plus maze surrounded by a square track (Fig. 1A). Liquid reward dispensers were located at each corner. The rat’s access to maze segments was controlled by 12 automatic doors located at maze intersections that could be raised and lowered vertically. The maze and surroundings (walls, floor, ceiling) were uniformly black with no landmarks to disambiguate the four cardinal directions. Four identical computer monitors were positioned 1 m above the four T-intersections on each side of the maze. All four monitors were continuously backlit throughout all experimental sessions; although the environment was dark, the dimly illuminated monitors were visible against the surrounding black curtains from most positions on the maze, forming a configuration of distal visual cues. When a distinguishing visual landmark was displayed (a white triangle with black center on one of the four monitors), the symmetry of the monitor array was broken, creating an asymmetric cue constellation that provided directional information. When no landmark was displayed, all four monitors appeared identical; the environment remained near-dark but the faintly visible monitors provided optical flow—useful for path integration computations—without directional landmark information.

To minimize influences of uncontrolled intramaze cues, 12 identical linear segments, 4 three-way intersections, and the center four-way intersection of the maze track were removed, cleaned, and replaced in a new random configuration prior to each day’s session. To minimize influences of uncontrolled extramaze cues, the task reference frame (start, goal, visual landmark) was rigidly rotated with respect to the room reference frame by a random multiple of 90° prior to each day’s session (and sometimes between trials within a session; see below). During behavior sessions the rat was illuminated by four IR lights located in the upper corners of the room.

During behavioral sessions, rats were monitored remotely from a control room using a custom interface developed in PyQt4. Video data, including real-time position information, were streamed via ZeroMQ from a Raspberry Pi 4 equipped with a Pi Camera HD (IR filter removed) mounted above the maze. Markerless motion tracking was performed directly on the Raspberry Pi using OpenCV, enabling lightweight, on-device processing of animal position without the need for physical markers.

Maze configurations (including changes to barrier positions and cue presentations) were coordinated through the control system using a MODBUS network for actuators and syringe pumps, and ZeroMQ for visual cue delivery. Barriers were raised and lowered by Progressive Automations linear actuators (PA-15-12-11), driven by BTS7960 motor drivers controlled by LAFVIN Nano V3.0 microcontrollers. These microcontrollers received MODBUS commands via TTL/RS485 two-way signal converters. Visual cues were displayed on monitors controlled by additional Raspberry Pi units running Python with the Pygame library, with control signals exchanged over Ethernet using ZeroMQ.

Additionally, to ensure that external noise did not disrupt behavior white noise was played in frequency range of 50hz to 20khz using a high frequency capable speaker. Noise generation was conducted using the python package PyAudio. Furthermore, a Hidizs DH805 digital to analog converter was used to ensure the full frequency range of sound was passed to the speaker. White noise was played during all sessions at an approximate 60db measured at corners of maze.

Finally, four custom designed syringe pumps utilizing nema 17 stepper motors, driven by TB67S249FTG stepper drivers and controlled by LAFVIN Nano V3.0 micro controllers that received MODBUS commands via TTL/RS485 two-way signal converters, were used to deliver liquid reward. The syringe pumps were able to deliver 100μl reward rapidly and consistently (±5μl) of sweetened condensed milk (SCM) mixed with water at a ratio 2 to 3 respectively. Syringe pumps were placed vertically under reward arms and fitted with 30ml syringes, connected to reward wells via Tygon tubing. Syringes were cleaned daily with water until they no longer held a proper seal and reward delivery was inconsistent. Lines running from the syringe pump were also flushed three times with water, 75% ethanol, and air after all the days sessions were completed.

### Exposure Sessions

After handling and reduction to 75% ad lib weight per protocol, rats were exposed to the maze in a series of two sessions. During the first exposure all barriers were in the down position for the entirety of the session and the rats were placed on a long arm segment closest to the entry point of the maze. Rats were then free to roam the maze. Upon entry to the first reward well they encountered, SCM was given. Each reward was followed by a pause during which no reward was available; the length of each pause was drawn from a Gaussian distribution (mean 45 s, SD 10 s). After the pause, reward was again available at any of the corner wells except the one that had most recently delivered reward. This continued until a total of 32 rewards were given or the session reached 30 minutes.

On the second day of exposure the barriers were raised at both ends of each linear arm segment surrounding the center of the maze. The rats started the session from the center of the maze unable to move to any other segment because again all central barriers were raised. After 2 minutes the east central barrier lowered. The rat then needed to enter the east arm and after another minute the north barrier would lower and the rat would then need to enter the north arm. This process continued in the counter clockwise direction until all central barriers were down. After entering the south arm and a minute had passed the barriers at the end of the central linear segments were then lowered and the rat was then free to search for reward in the same manner as the first session. The session ended when the rat consumed 32 rewards or the session reached 30 minutes.

### Pseudo Random Trial Generation

Trial sequences were pseudorandomly generated in shuffled blocks of eight. Sequences were constrained such that runs of three consecutive trials from the same start arm occurred no more than twice per session and, when occurring twice, were distributed across the first and second halves of the session. Equivalent constraints were applied to route and goal sequences in the VC protocol.

### Statistical Analysis

Analyses were performed in Python 3.12 (pandas 3.0, SciPy 1.17, statsmodels 0.14, pingouin 0.6, scikit-learn 1.8) and R 4.5 (afex 1.5, called from Python through rpy2). Unless stated otherwise, data are reported as mean ± SEM, tests were two-tailed, and effects were considered significant at α = .05. The acquisition criterion (at least 24 correct of 32 trials in a session), the reversal criterion (8 correct trials within a block of 8) and the above-chance thresholds used for the probe tests were derived from the binomial distribution, assuming two equiprobable response alternatives (p = .5 for trials from trained start arms and p = .25 for Novel Route trials) and a chance probability of ≤ .01.

Repeated-measures designs, with trial block or session as the within-subjects factor and training group as the between-subjects factor, were analyzed with mixed ANOVAs using Type III sums of squares (afex). Sphericity was assessed with Mauchly’s test for every within-subjects factor with more than two levels; where it was violated (p < .05), Greenhouse–Geisser-corrected degrees of freedom and p values are reported and the correction is noted in the text. Post hoc pairwise comparisons following a significant effect were Bonferroni-corrected t tests (Welch-corrected where group sizes were unequal). Planned comparisons between pooled groups were tested with single-degree-of-freedom contrasts on the ANOVA error term or with Welch t tests, as indicated.

Sessions and trials to criterion were compared across groups with Kruskal–Wallis tests followed, where indicated, by Holm-corrected Mann–Whitney U tests, or with one-way ANOVA. For the reversal probe, rats removed from the maze after reaching criterion were credited with 100% accuracy for the remaining trial blocks, and the two rats whose sessions ended before they had reached criterion were excluded from all reversal analyses. Trials whose route could not be classified were excluded from the perseveration analysis (see Results).

For the individual-differences analysis (Fig. 5), each PI+VC rat’s Novel Route, No-Cue and Rotation scores (proportion of correct trials) were z-scored and submitted to principal component analysis; because the first two components explained 90.2% of the variance, k-means clustering (k = 3, 50 random initializations) was performed in the two-dimensional component space. The number of clusters was selected with the gap statistic (500 uniform reference datasets) and the Calinski–Harabasz index, and the stability of the solution was assessed by bootstrap resampling (1,000 resamples, k-means refit in the fixed component space with all rats reassigned), cluster-wise Jaccard similarity (Hennig, 2007), leave-one-out linear discriminant classification and leave-one-rat-out reclustering. Reversal performance was withheld from clustering and scored as rescaled trials to criterion (see Results); cluster differences were tested with one-way ANOVA, accompanied by Levene’s test, Welch’s ANOVA and the Kruskal–Wallis test, and with a log-rank test on trials to criterion in which rats that never reached criterion were treated as censored. The relationship between probe and reversal performance was assessed with multiple regression using HC3 heteroscedasticity-robust standard errors and with Spearman rank correlations, and associations between cluster membership and sex or cue-approach direction with χ² tests.

### Use of Artificial Intelligence

Claude (Anthropic, via Claude Code, 2025–2026) assisted with writing the analysis code for data processing, statistics, figures and supplementary-video annotation; Claude, ChatGPT (OpenAI) and Undermind.ai assisted with literature searches. The authors reviewed and tested all AI-assisted code, verified every reported statistic against the data, and made all design, analytical and interpretive decisions.

### Supporting Information

**Video S1. Basic structure of the 2S2C task.**

Example trials from a PI+VC rat on its fifth acquisition session: placement on the maze, two consecutive trials from the proximal and distal start arms with the return to the start arm between them, and four consecutive correct trials from a VC rat with the landmark in four different positions. The PI condition uses the identical trial structure with the monitors dimly backlit. Real time; the overlay shows trial number and running score. MP4 (H.264), 728 × 728 pixels, 30 fps, 2 min 59 s.

**Video S2. Deliberation on the first Novel Route trial.**

A familiar-route trial, the first Novel Route trial (trial 17) and the last Novel Route trial (trial 40) from one PI_PI_ rat and one VC_VC_ rat. On trial 17 the PI-trained rat shuttles between the novel start arm and the maze center for more than 30 s before choosing, whereas the VC-trained rat runs without pausing to the well that the trained turn sequence would have led to. By trial 40 the PI-trained rat takes the direct novel route while the VC-trained rat still repeats the trained sequence. Clips end at the first well visited. MP4 (H.264), 728 × 728 pixels, 30 fps, 2 min 12 s.

**Video S3. Perseveration during the Reversal probe.**

A pre-reversal familiar trial, the first Reversal trial (trial 17) and the last trial of the session from one PI_PI_ rat and one VC^VC^ rat, each shown in full; the asterisk marks the current goal. On trial 17 both rats run first to the former goal. By its last trial (trial 40) the PI_PI_ rat, having reached criterion, runs directly to the new goal, whereas the VC^VC^ rat still visits the former goal first on its 80th trial and never reached criterion. MP4 (H.264), 728 × 728 pixels, 30 fps, 4 min 13 s.

## Acknowledgements

This work was supported by NIH U01NS128664 and NIH U01NS126050 to H.T.B. We thank Alicia Izquierdo, Aaron Blaisdell, and Andrew Wikenheiser for helpful comments and discussion. ChatGPT (OpenAI) and Claude (Anthropic) were used to revise and edit portions of the text. The authors reviewed all text and take full responsibility for the content.

